# The Longitudinal Assessment of Neuropsychiatric Symptoms in Mild Cognitive Impairment and Alzheimer’s disease and their Association with White Matter Hyperintensities in the National Alzheimer’s Coordinating Center’s Uniform Data Set

**DOI:** 10.1101/822809

**Authors:** Cassandra J. Anor, Mahsa Dadar, D. Louis Collins, M. Carmela Tartaglia

**Author notes:** These authors contributed equally to this work (Cassandra J. Anor, Mahsa Dadar). Correspondence to: Carmela Tartaglia, M.D., FRCPC, Marion and Gerald Soloway Chair in Brain Injury and Concussion Research, Associate Professor, Tanz Centre for Research in Neurodegenerative Diseases, University of Toronto, Cognitive Neurologist, Memory Clinic - Toronto Western Hospital, Director Memory Clinical Trials Unit, 399 Bathurst St. WW 5-449, Toronto, ON M5T 2S8. **FINANCIAL DISCLOSURES:** None of the authors have any financial disclosures.

## Abstract

**Introduction:** Neuropsychiatric symptoms (NPS) are common in all dementias, including those with Alzheimer’s disease (AD). NPS contribute to patients’ distress, caregiver burden, and can lead to institutionalization. White matter hyperintensities (WMH) are a common finding on MRI usually indicative of cerebrovascular disease and have been associated with certain NPS. The aim of this study was two-fold. Firstly, we assessed the relationship between WMH load and NPS severity in MCI due to AD (MCI-AD) and AD. Secondly, we assessed the ability of WMH to predict the development and progression of NPS in these participants. Data was obtained from the National Alzheimer’s Coordinating Center.

**Methods:** WMH were obtained from baseline MRIs and quantified using an automated segmentation technique. NPS were measured using the Neuropsychiatric Inventory (NPI). Mixed effect models and correlations were used to determine the relationship between WMH load and NPS severity scores.

**Results:** Cross-sectional analysis showed no significant association between NPS and WMH at baseline. Longitudinal mixed effect models, however, revealed a significant relationship between increase in NPI total scores and baseline WMH load (p=0.014). There was also a significant relationship between increase in irritability severity scores over time and baseline WMH load (p= 0.009). Trends were observed for a relationship between increase in agitation severity scores and baseline WMH load (p=0.058). No other NPS severity scores were significantly associated with baseline WMH load. The correlation plot analysis showed that baseline whole brain WMH predicted change in future NPI total scores (r=0.169, p=0.008). Baseline whole brain WMH also predicted change in future agitation severity scores (r= 0.165, p= 0.009). The temporal lobe WMH (r=0.169, p=0.008) and frontal lobe WMH (r=0.153, p=0.016) contributed most to this this change.

**Conclusion:** Irritability and agitation are common NPS and very distressful to patients and caregivers. Our findings of an increase in irritability severity over time as well as higher agitation severity scores at follow-up in participants with MCI-AD and AD with increased WMH loads have important implications for treatment, arguing for aggressive treatment of vascular risk factors in patients with MCI-AD and AD.

## 1. INTRODUCTION

Neuropsychiatric symptoms (NPS) are common in patients with dementia and are a major source of patients’ distress, caregiver burden, and can lead to institutionalization(Huang et al., 2012; Steele et al., 1990). Moreover, while psychotropic drugs may temporarily alleviate certain NPS, some have severe and harmful effects(I. and J., 2005). As such, there is a need to better understand NPS and their underlying pathology. In patients with Alzheimer’s disease (AD), the most commonly reported NPS are apathy, agitation(Koenig et al., 2016; Q.-F. et al., 2016), irritability(Koenig et al., 2016), and depression(Boublay et al., 2016; Q.-F. et al., 2016). Some studies have reported the presence and severity of depressive symptoms as predictive of MCI progression to AD(M. et al., 2008; Van Der Mussele et al., 2014), but this has not been observed in all studies(Palmer et al., 2010, 2007)

The cause of NPS in dementia is unknown, but there is some evidence that focal white matter hyperintensities (WMH) are associated with NPS(Kim et al., 2013; Pantoni, 2010). WMH are areas of increased signal relative to the surrounding white matter regions, as seen on magnetic resonance imaging (MRI) T2-weighted and fluid attenuated inversion recovery (FLAIR) sequences(F. et al., 2018). WMH can be seen in cognitively normal individuals and are known to increase with age. The pathological correlates of WMH are heterogeneous including edema, inflammation or infection, but are most commonly indicative of cerebral small vessel disease (SVD)(Pantoni, 2010). Cerebral SVD is associated with vascular risk factors, such as hypertension, hypercholesterolemia, and diabetes. Some studies have shown an association between increased WMH and AD(Erkinjuntti et al., 1994; Leys et al., 1990; Yoshita et al., 2006). In a recent systematic review, Liu and colleagues (2018) found that cerebral SVD markers, such as WMHs and cerebral micro-bleeds, were associated with increased risk of clinically diagnosed AD(Y. et al., 2018).

WMH have been associated with specific NPS in patients with dementia. In AD, depression(Clark et al., 1998), aberrant motor behaviours(Berlow et al., 2010; N. Hirono et al., 2000; Nobutsugu Hirono et al., 2000), anxiety(Berlow et al., 2010), night-time behaviours(Berlow et al., 2010), and apathy(Starkstein et al., 1997) have been reported in association with increased prevalence of WMH. Some studies have related periventricular WMH volumes, especially in the frontal lobes, to hallucinations, depression, and anxiety in patients with AD and Vascular dementia (VaD)(Park et al., 2011; Soennesyn et al., 2012; Starkstein et al., 2009). Focal frontal WMH have also been associated with depression in patients with dementia with Lewy bodies(Barber et al., 1999), apathy in vascular cognitive impairment(Kim et al., 2013), verbal aggressiveness in amnestic MCI and AD(Ogama et al., 2018), and delusions in patients with AD(Cassandra J. Anor et al., 2017; M. and M., 2015; Ogawa et al., 2013). Parieto-occipital white matter changes have been shown to contribute to the development of delusional misidentification in patients with AD(Lee et al., 2006). However, there are also reports of no significant relationship between NPS and WMH in patients with AD(Klugman et al., 2009; Modrego et al., 2008). Given the inconclusiveness of the current literature, there is a need to better understand the relationship between WMH and NPS in patients with MCI or AD.

The present study explores the relationship between WMH and NPS in patients with MCI-AD and AD from a large cohort of patients collected by the National Alzheimer’s Coordinating Center (NACC). Our study aims were to determine the relationship between NPS and WMH burden in patients with MCI-AD or AD and assess the relationship between WMH volume and development of specific NPS in a longitudinal cohort of these patients. A clear relationship between NPS and WMH may justify more aggressive treatment of vascular risk factors given the implication of NPS in patients’ distress, caregiver burden, and institutionalization.

## 2. MATERIALS AND METHODS

### 2.1 PARTICIPANTS

We used data collected by the National Alzheimer’s Coordinating Center’s Uniform Data Set (NACC), n=753. NACC developed a database of standardized clinical research data obtained from 34 past and present NIA-funded Alzheimer’s Disease Centers across the United States. NACC data collection has been previously described elsewhere(Beekly et al., 2004; Morris et al., 2006). Participants included in the database have a range of cognitive statuses: normal cognition, mild cognitive impairment (MCI), and demented due to various clinical diagnoses.

Participants in our study had to have a diagnosis of MCI due to AD (MCI-AD) or AD in accordance with NINCDS-ADRDA criteria(McKhann et al., 1984), aged 50 or older, a Neuropsychiatric Inventory (NPI) and a baseline MRI. We removed participants whose clinical data and MR imaging data were not within three months of each other and those who did not have at least one follow up clinic visit (N=447). Other exclusion criteria included: history of seizures, psychiatric disorders, traumatic brain injury, stroke, other cognitive disorders, and primary etiologic diagnoses other than AD. The final study cohort consisted of 252 participants (N=114 with MCI-AD, N=138 with AD).

### 2.2 VARIABLES

The following NACC data was used for this study: diagnosis, MRI data, age, education, clinical dementia rating (CDR) sum of boxes scores, presence/absence of vascular risk factors presence/absence and severity of NPS, and the use of psychotropic and/or vascular risk factor-controlling medications. A vascular risk factor included: history of/or currently smoking, heart attack/cardiac arrest, atrial fibrillation, angioplasty/ endarterectomy/ stent, bypass surgery, pacemaker, congestive heart failure, other cardiovascular disease, TIAs, diabetes, hypertension, hypercholesterolemia, and whether the subject was currently taking any vascular medications. Medication to control vascular risk factors included the use of the following categories: anti-adrenergic, anticoagulant/antiplatelet, ACE inhibitor, antihypertensive or blood pressure medication, angiotensin II inhibitor, beta blocker, calcium channel blocking agent, diabetes medication, diuretic, antihypertensive combination, lower lipid levels, and the use of a vasodilator.

The NPI was used to evaluate NPS. The NPI is based on scripted questions administered to the participants’ caregivers or an informant familiar with the participant(Cummings et al., 1994). It is used to evaluate the presence, severity, and frequency of twelve commonly encountered NPS in dementia: delusions, hallucinations, agitation, depression, anxiety, elation, apathy or indifference, disinhibition, irritability, aberrant motor behaviour, night-time behaviour, and appetite or eating(Cummings et al., 1994). Furthermore, the NPI is a commonly used scale to assess patients with dementias and other neurological disorders with acceptable validity and reliability in outpatient settings(Cummings et al., 1994). We focused on the presence or absence and the severity of each symptom. The severity of each symptom is scored on a 3-point scale (1=mild, 2=moderate, 3=severe).

Since it is well known that medications can have an impact on NPS, we took psychotropic medication usage into account. Data on psychotropic medication use was provided by NACC, specifically the use of antipsychotics, antidepressants, and anxiolytics. Antipsychotics, antidepressants, and anxiolytics were coded as presence/absence (0/1). We added the variable presence/absence (1/0) of sleep medications. A coding of 1 was given if any of the following sleep medications were listed: Ambien, melatonin, Zolpidem, and/or Zopiclone. The presence/absence of these psychotropic medications were included in our correlation plots of baseline whole brain WMH load, used to predict that change in future NPI severity scores over time.

### 2.3 MRI PROCESSING

NACC participants diagnosed with either MCI-AD or AD, with a baseline MRI that included a FLAIR sequence within 3 months of clinical assessment were used for analysis in our study. Only baseline MRIs were used for this study, as we did not have enough follow up scans to match follow up NPI scores. The MRI parameters have been previously described elsewhere(M. Dadar et al., 2017). All MRI scans were pre-processed in three steps: i) denoising(Manjón et al., 2010), ii) intensity inhomogeneity correction(Zijdenbos and Evans, 1998), and iii) intensity normalization(Fonov et al., 2011). FLAIR scans were rigidly co-registered to the T1W scans(Collins et al., 1994). The T1W scans were registered to the ADNI template(Collins and Evans, 1997). Concatenating the two transformations, the FLAIR scans were also registered to the ADNI template. Using a previously validated automatic WMH segmentation technique and a library of manual segmentations from the NACC dataset, the WMHs were segmented automatically(Dadar et al., 2018; M. Dadar et al., 2017; Mahsa Dadar et al., 2017). The quality of the segmentations was assessed and verified by an expert (MD). The total volumes of the WMHs were then calculated (in mm^3^) and normalized for head size and log-transformed to achieve normal distribution.

### 2.4 STATISTICAL ANALYSIS

Cross sectional analysis was performed in order to assess the association between baseline total NPS severity scores and baseline whole brain WMH load. WMH load, age, and education were used as continuous predictors. Sex was used as a categorical predictor In addition, the cross sectional analysis was repeated for the twelve individual NPI severity scores. FDR was used to correct for multiple comparisons.

Longitudinal mixed-effects models were used to assess the association of WMH load at baseline with the longitudinal changes in NPI symptoms. WMH load, age, education, as well as the baseline NPI scores were used as continuous predictors. Sex was used as a categorical predictor. This analysis was repeated for the twelve individual NPI subscores, but using the respective baseline subscore as a continuous predictor. FDR was used to correct for multiple comparisons.

Correlation plots were used to determine if baseline whole brain WMH load could predict change in future NPI severity scores. Subjects were considered as categorical random effects in all the mixed effects models. WMH volumes were log-transformed to achieve normal distribution. All continuous variables were z-scored prior to the analysis. Mixed effects models were fitted using fitlme in MATLAB version R2017b. P-values reported are not corrected for multiple comparisons.

For any sub-score correlations that are statistically significant, we completed a secondary analysis to investigate which lobe contributed to the correlation.

## 3. RESULTS

The final cohort consisted of 252 participants (138AD, 114MCI-AD), 134 were male, 118 were female. The average number of years of education was 15.65±8.35 (Table 1). There was an average of 22.8 days in between the baseline NPI and the baseline MRI scan. The usage of psychotropic medications and medications to control vascular risk factors were reported at baseline (Table 2). Each of the clinic visits thereafter, which included NPI scores, were approximately one year after the baseline MRI scan (see Appendix, Supplementary Table 1). Whole brain and lobar WMH volumes were calculated for the baseline MRI scans (Table 3). At baseline, all twelve NPS were present (see Appendix, Supplementary Table 2). All twelve NPS were also present at each follow-up visit (Table 4).

**Table 1.** Basic demographics at baseline visit

| Diagnosis | MCI-AD | AD |
| --- | --- | --- |
| N | 114 | 138 |
| Age (mean ± SD) | 76.1 ± 7.12 | 75.8 ± 8.62 |
| Sex (M/F) | 68/46 | 66/72 |
| Education (mean ±SD) | 16.4 ± 8.54 | 15.0 ± 8.16 |

**Table 2.**
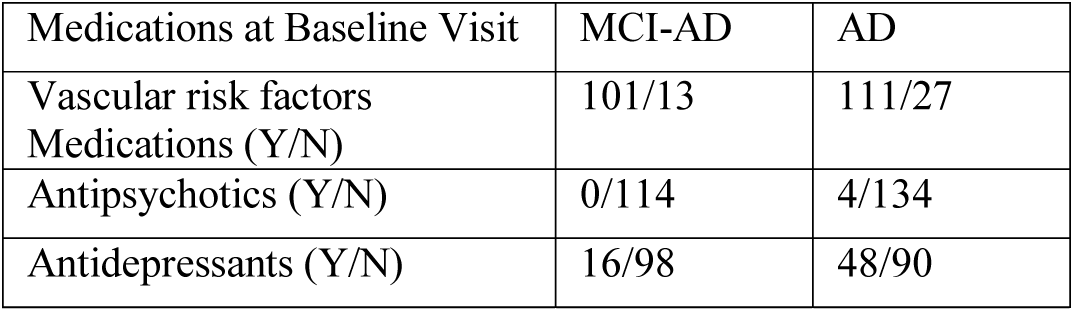
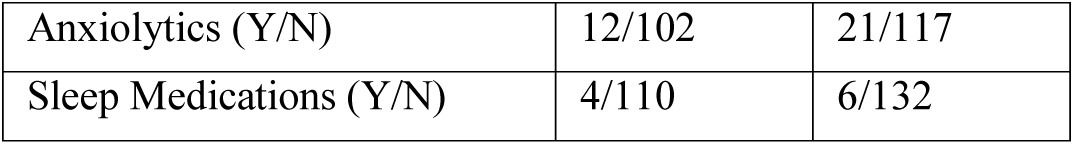
Absence/presence (N/Y) of various medications at baseline visit

| Medications at Baseline Visit | MCI-AD | AD |
| --- | --- | --- |
| Vascular risk factors Medications (Y/N) | 101/13 | 111/27 |
| Antipsychotics (Y/N) | 0/114 | 4/134 |
| Antidepressants (Y/N) | 16/98 | 48/90 |
| Anxiolytics (Y/N) | 12/102 | 21/117 |
| Sleep Medications (Y/N) | 4/110 | 6/132 |

**Table 3.**
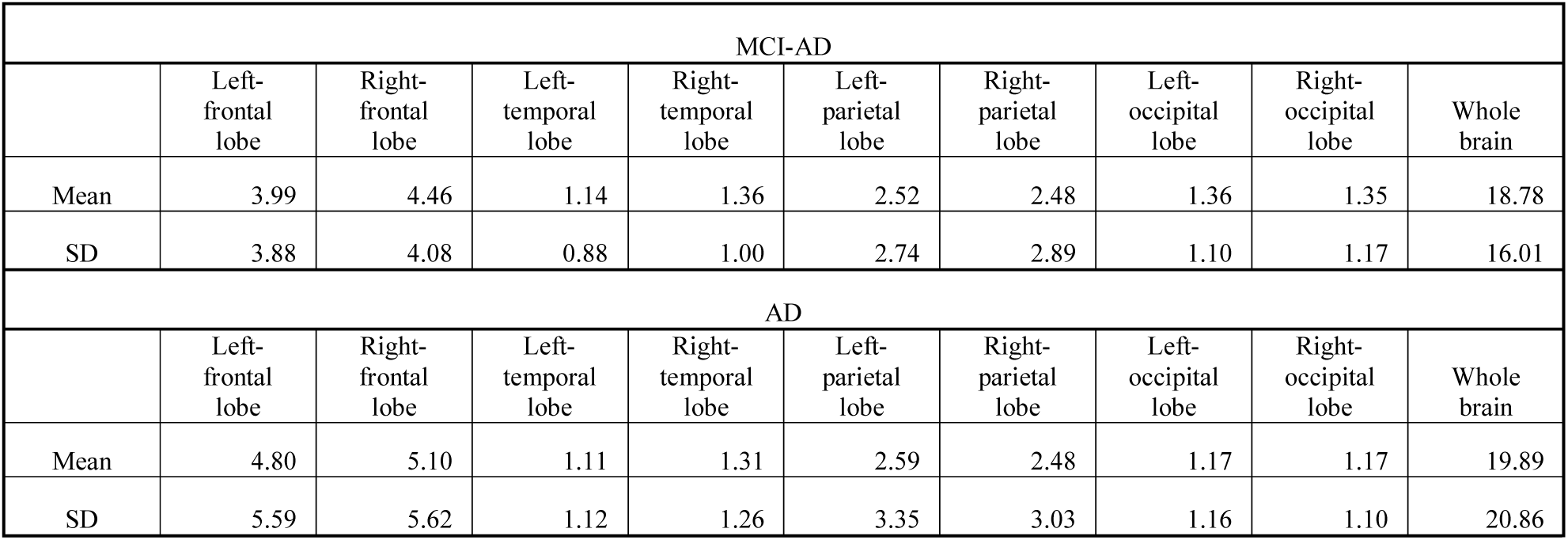
Regional WMH volumes (mm^3^) for baseline MRI scan

| MCI-AD |  |  |  |  |  |  |  |  |  |
| --- | --- | --- | --- | --- | --- | --- | --- | --- | --- |
|  | Left-frontal lobe | Right-frontal lobe | Left-temporal lobe | Right-temporal lobe | Left-parietal lobe | Right-parietal lobe | Left-occipital lobe | Right-occipital lobe | Whole brain |
| Mean | 3.99 | 4.46 | 1.14 | 1.36 | 2.52 | 2.48 | 1.36 | 1.35 | 18.78 |
| SD | 3.88 | 4.08 | 0.88 | 1.00 | 2.74 | 2.89 | 1.10 | 1.17 | 16.01 |
| AD |  |  |  |  |  |  |  |  |  |
|  | Left-frontal lobe | Right-frontal lobe | Left-temporal lobe | Right-temporal lobe | Left-parietal lobe | Right-parietal lobe | Left-occipital lobe | Right-occipital lobe | Whole brain |
| Mean | 4.80 | 5.10 | 1.11 | 1.31 | 2.59 | 2.48 | 1.17 | 1.17 | 19.89 |
| SD | 5.59 | 5.62 | 1.12 | 1.26 | 3.35 | 3.03 | 1.16 | 1.10 | 20.86 |

**Table 4.** Prevalence (%) of NPS sub-score at each clinical visit

|  | NPS (%) |  |  |  |  |  |
| --- | --- | --- | --- | --- | --- | --- |
| Visit Number | Visit 1<br>(N=252) | Visit 2<br>(N=247) | Visit 3<br>(N=184) | Visit 4<br>(N=117) | Visit 5<br>(N=55) | Visit 6<br>(N=30) |
| Delusions | 7.1 | 12.6 | 13.6 | 15.4 | 10.9 | 20.0 |
| Hallucinations | 2.0 | 6.9 | 7.6 | 8.5 | 7.3 | 6.7 |
| Agitation | 27.0 | 32.0 | 38.0 | 23.9 | 40.0 | 16.7 |
| Depression | 29.8 | 34.8 | 34.8 | 27.4 | 21.8 | 33.3 |
| Anxiety | 30.2 | 32.0 | 35.9 | 40.2 | 29.1 | 36.7 |
| Elation | 2.8 | 6.5 | 4.3 | 6.0 | 1.8 | 3.3 |
| Apathy | 31.0 | 34.4 | 41.8 | 44.4 | 45.5 | 46.7 |
| Disinhibition | 17.5 | 19.8 | 15.8 | 20.5 | 27.3 | 26.7 |
| Irritability | 36.1 | 36.0 | 31.5 | 36.8 | 21.8 | 40.0 |
| Aberrant Motor behaviour | 14.7 | 20.2 | 25.5 | 23.1 | 16.4 | 33.3 |
| Night-time Behaviour | 27.0 | 30.4 | 28.3 | 33.3 | 36.4 | 30.0 |
| Appetite | 17.1 | 27.1 | 26.6 | 27.4 | 30.9 | 33.3 |

The cross-sectional analysis at baseline revealed no significant association between any of the NPS scores (either total or subscores) and WMH (see Appendix, Supplementary Table 2).

With regards to the longitudinal mixed effect model analyses, there was a statistically significant relationship (after correction for multiple comparisons) between change in NPI total scores and baseline whole brain WMH load (p= 0.014) (Table 5). For the longitudinal mixed effect model analyses of the NPS subscores, there was a statistically significant relationship between the longitudinal increase in the irritability severity score and baseline WMH load (p= 0.009). There were also trends for an increase in agitation severity score (p=0.058) and elation severity score and baseline WMH load (p= 0.061). However, the trend for the increased elation score is based on a few subjects who reported the presence of elation (only 2.8-6.5% of the sample cohort at each clinic visit). No other NPS severity scores were significantly associated with baseline WMH load (Table 6).

**Table 5.**
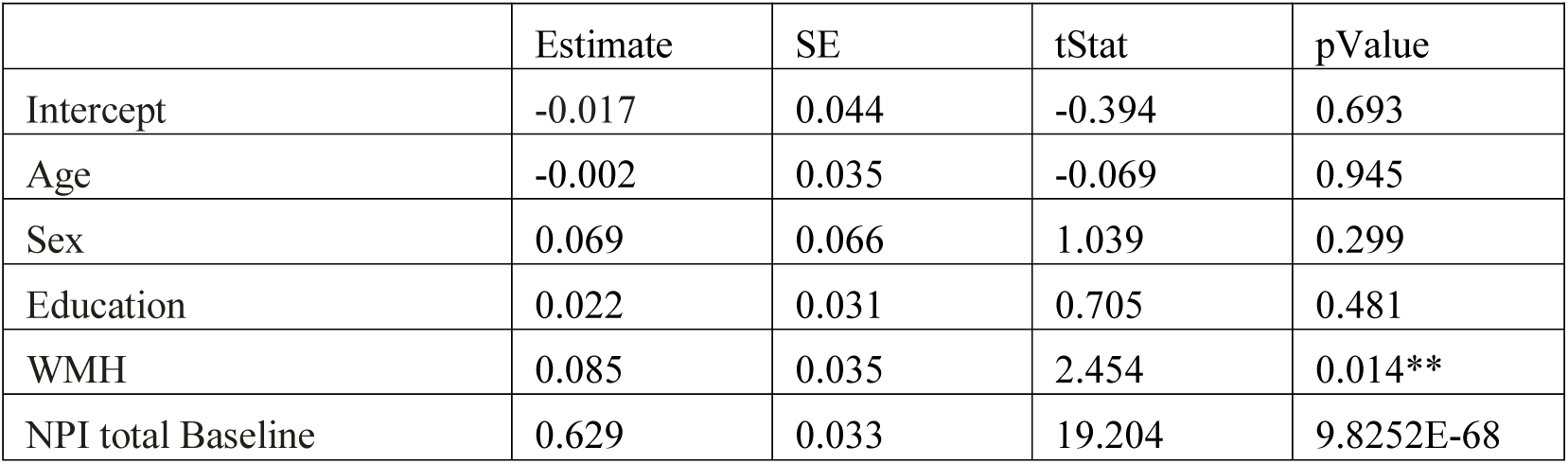
Longitudinal Mixed-Effect Models of annualized change in NPI total score explained by whole brain WMH load, age, sex, education and baseline total NPI scores. ** denotes statistical significance, ◊Denotes a trend; one-tailed test p<0.1

|  | Estimate | SE | tStat | pValue |
| --- | --- | --- | --- | --- |
| Intercept | -0.017 | 0.044 | -0.394 | 0.693 |
| Age | -0.002 | 0.035 | -0.069 | 0.945 |
| Sex | 0.069 | 0.066 | 1.039 | 0.299 |
| Education | 0.022 | 0.031 | 0.705 | 0.481 |
| WMH | 0.085 | 0.035 | 2.454 | 0.014** |
| NPI total Baseline | 0.629 | 0.033 | 19.204 | 9.8252E-68 |

**Table 6.**
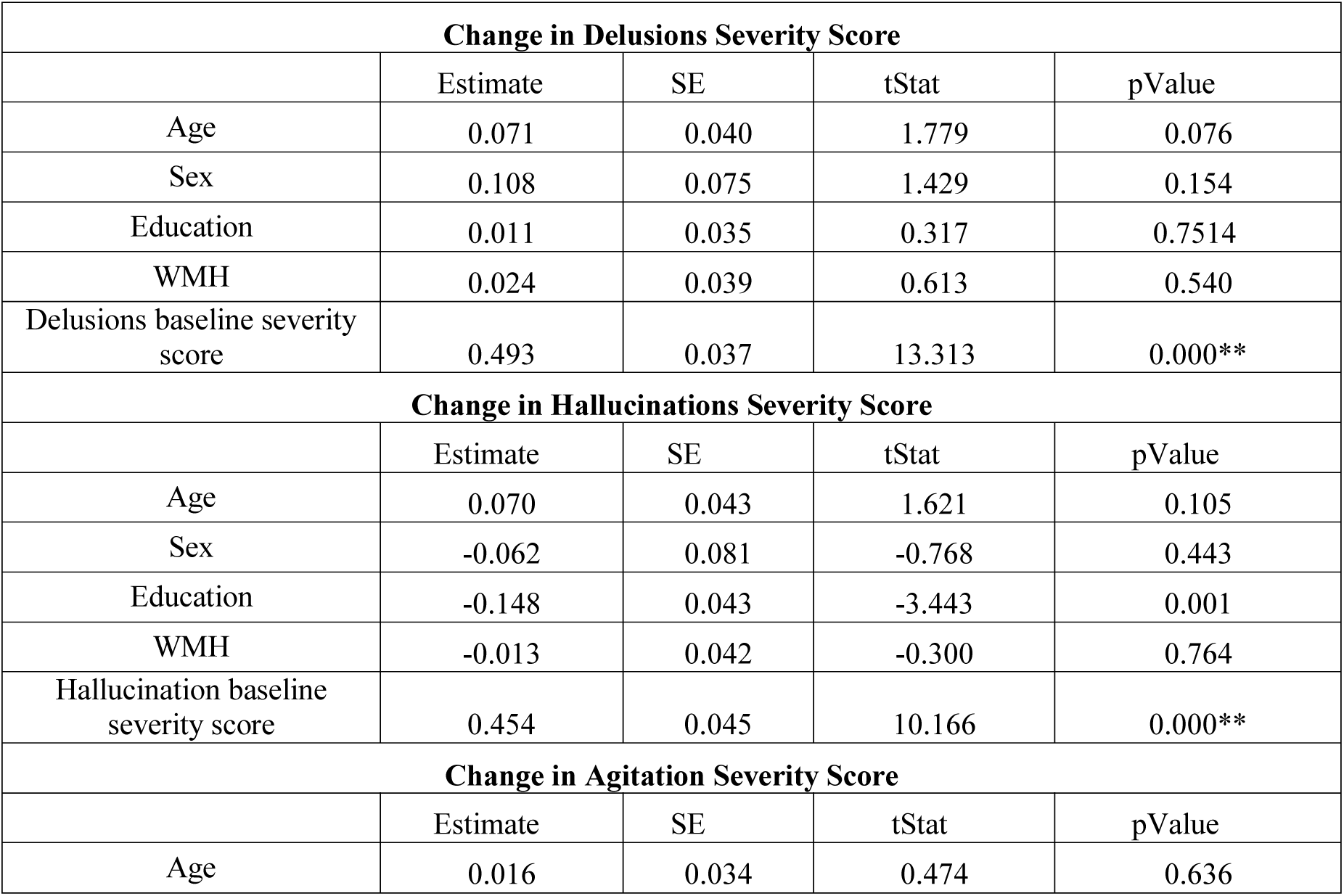
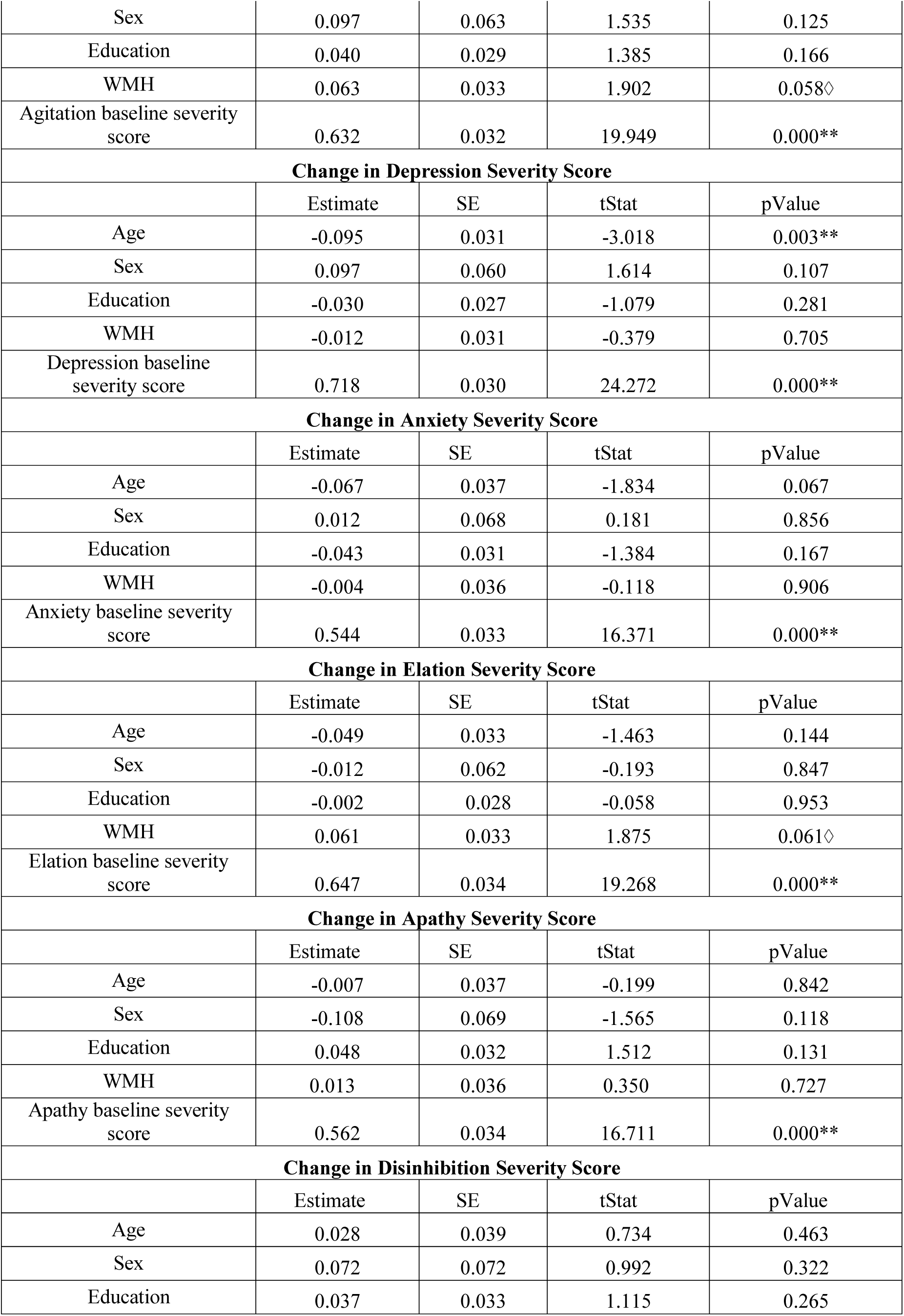
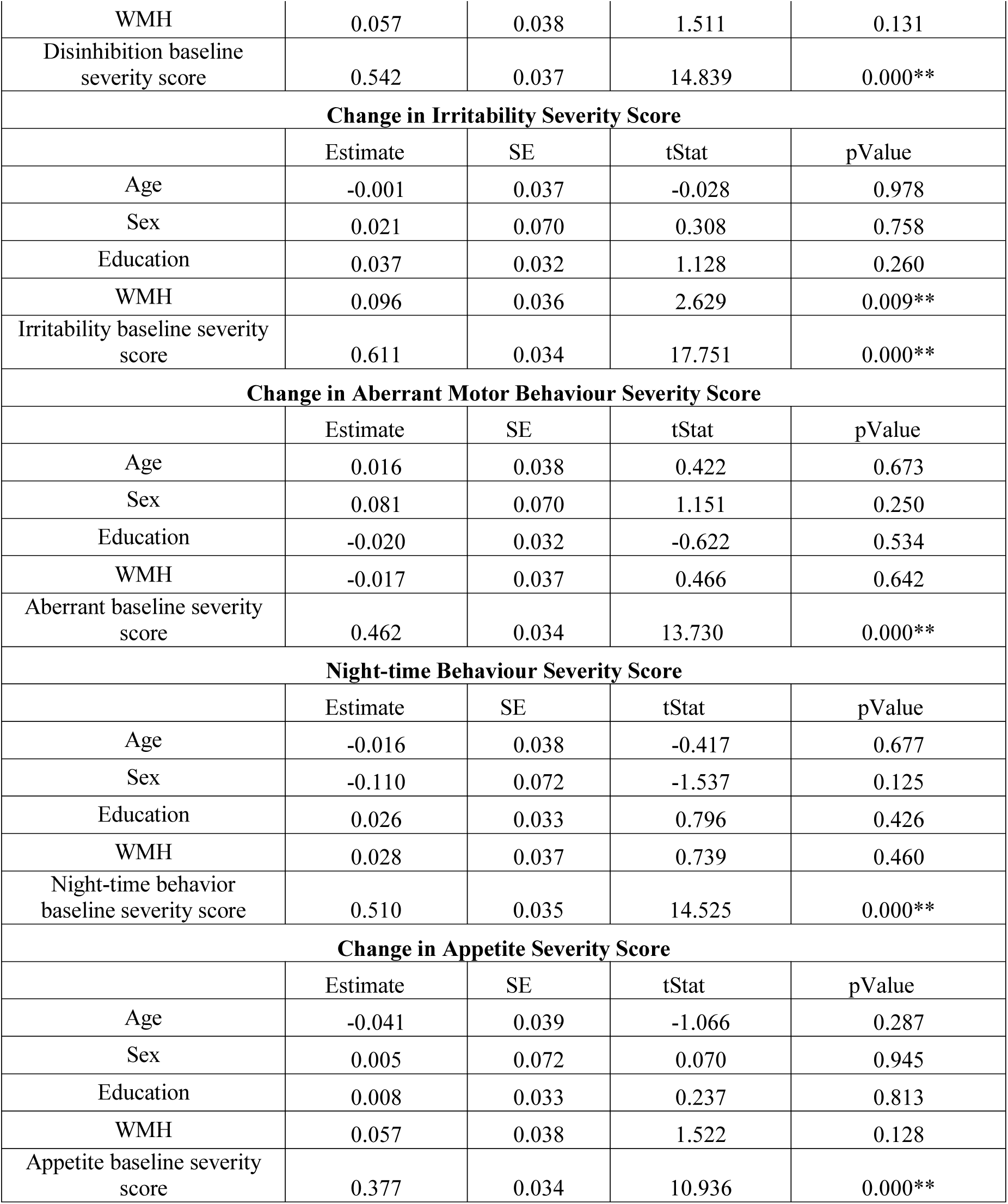
Longitudinal Mixed-Effect Models of Annualized Change in NPI subscores explained by age, sex, education, whole brain WMH load and NPS severity scores. ** denotes statistical significance, ◊Denotes a trend; one-tailed test p<0.1

| **Change in Delusions Severity Score** |  |  |  |  |
| --- | --- | --- | --- | --- |
|  | Estimate | SE | tStat | pValue |
| Age | 0.071 | 0.040 | 1.779 | 0.076 |
| Sex | 0.108 | 0.075 | 1.429 | 0.154 |
| Education | 0.011 | 0.035 | 0.317 | 0.7514 |
| WMH | 0.024 | 0.039 | 0.613 | 0.540 |
| Delusions baseline severity score | 0.493 | 0.037 | 13.313 | 0.000** |
| **Change in Hallucinations Severity Score** |  |  |  |  |
|  | Estimate | SE | tStat | pValue |
| Age | 0.070 | 0.043 | 1.621 | 0.105 |
| Sex | -0.062 | 0.081 | -0.768 | 0.443 |
| Education | -0.148 | 0.043 | -3.443 | 0.001 |
| WMH | -0.013 | 0.042 | -0.300 | 0.764 |
| Hallucination baseline severity score | 0.454 | 0.045 | 10.166 | 0.000** |
| **Change in Agitation Severity Score** |  |  |  |  |
|  | Estimate | SE | tStat | pValue |
| Age | 0.016 | 0.034 | 0.474 | 0.636 |
| Sex | 0.097 | 0.063 | 1.535 | 0.125 |
| Education | 0.040 | 0.029 | 1.385 | 0.166 |
| WMH | 0.063 | 0.033 | 1.902 | 0.058◇ |
| Agitation baseline severity score | 0.632 | 0.032 | 19.949 | 0.000** |
| **Change in Depression Severity Score** |  |  |  |  |
|  | Estimate | SE | tStat | pValue |
| Age | -0.095 | 0.031 | -3.018 | 0.003** |
| Sex | 0.097 | 0.060 | 1.614 | 0.107 |
| Education | -0.030 | 0.027 | -1.079 | 0.281 |
| WMH | -0.012 | 0.031 | -0.379 | 0.705 |
| Depression baseline severity score | 0.718 | 0.030 | 24.272 | 0.000** |
| **Change in Anxiety Severity Score** |  |  |  |  |
|  | Estimate | SE | tStat | pValue |
| Age | -0.067 | 0.037 | -1.834 | 0.067 |
| Sex | 0.012 | 0.068 | 0.181 | 0.856 |
| Education | -0.043 | 0.031 | -1.384 | 0.167 |
| WMH | -0.004 | 0.036 | -0.118 | 0.906 |
| Anxiety baseline severity score | 0.544 | 0.033 | 16.371 | 0.000** |
| **Change in Elation Severity Score** |  |  |  |  |
|  | Estimate | SE | tStat | pValue |
| Age | -0.049 | 0.033 | -1.463 | 0.144 |
| Sex | -0.012 | 0.062 | -0.193 | 0.847 |
| Education | -0.002 | 0.028 | -0.058 | 0.953 |
| WMH | 0.061 | 0.033 | 1.875 | 0.061◇ |
| Elation baseline severity score | 0.647 | 0.034 | 19.268 | 0.000** |
| **Change in Apathy Severity Score** |  |  |  |  |
|  | Estimate | SE | tStat | pValue |
| Age | -0.007 | 0.037 | -0.199 | 0.842 |
| Sex | -0.108 | 0.069 | -1.565 | 0.118 |
| Education | 0.048 | 0.032 | 1.512 | 0.131 |
| WMH | 0.013 | 0.036 | 0.350 | 0.727 |
| Apathy baseline severity score | 0.562 | 0.034 | 16.711 | 0.000** |
| **Change in Disinhibition Severity Score** |  |  |  |  |
|  | Estimate | SE | tStat | pValue |
| Age | 0.028 | 0.039 | 0.734 | 0.463 |
| Sex | 0.072 | 0.072 | 0.992 | 0.322 |
| Education | 0.037 | 0.033 | 1.115 | 0.265 |
| WMH | 0.057 | 0.038 | 1.511 | 0.131 |
| Disinhibition baseline severity score | 0.542 | 0.037 | 14.839 | 0.000** |
| **Change in Irritability Severity Score** |  |  |  |  |
|  | Estimate | SE | tStat | pValue |
| Age | -0.001 | 0.037 | -0.028 | 0.978 |
| Sex | 0.021 | 0.070 | 0.308 | 0.758 |
| Education | 0.037 | 0.032 | 1.128 | 0.260 |
| WMH | 0.096 | 0.036 | 2.629 | 0.009** |
| Irritability baseline severity score | 0.611 | 0.034 | 17.751 | 0.000** |
| **Change in Aberrant Motor Behaviour Severity Score** |  |  |  |  |
|  | Estimate | SE | tStat | pValue |
| Age | 0.016 | 0.038 | 0.422 | 0.673 |
| Sex | 0.081 | 0.070 | 1.151 | 0.250 |
| Education | -0.020 | 0.032 | -0.622 | 0.534 |
| WMH | -0.017 | 0.037 | 0.466 | 0.642 |
| Aberrant baseline severity score | 0.462 | 0.034 | 13.730 | 0.000** |
| **Night-time Behaviour Severity Score** |  |  |  |  |
|  | Estimate | SE | tStat | pValue |
| Age | -0.016 | 0.038 | -0.417 | 0.677 |
| Sex | -0.110 | 0.072 | -1.537 | 0.125 |
| Education | 0.026 | 0.033 | 0.796 | 0.426 |
| WMH | 0.028 | 0.037 | 0.739 | 0.460 |
| Night-time behavior baseline severity score | 0.510 | 0.035 | 14.525 | 0.000** |
| **Change in Appetite Severity Score** |  |  |  |  |
|  | Estimate | SE | tStat | pValue |
| Age | -0.041 | 0.039 | -1.066 | 0.287 |
| Sex | 0.005 | 0.072 | 0.070 | 0.945 |
| Education | 0.008 | 0.033 | 0.237 | 0.813 |
| WMH | 0.057 | 0.038 | 1.522 | 0.128 |
| Appetite baseline severity score | 0.377 | 0.034 | 10.936 | 0.000** |

In the correlation plots, baseline whole brain WMH load predicted the change in total NPI scores at a future visit (r=0.169, p=0.008) (Figure 1). Whole brain WMH load also predicted the change in some NPI severity sub-scores at a future visit. More specifically, there was a significant correlation between baseline WMH load and severity of agitation (r= 0.165, p= .009) (Figure 2). This relationship withstood correction for multiple comparisons and controlling for age, sex, and education. There was also a correlation between baseline whole brain WMH load and severity of irritability that although statistically significant, did not withstand correction for multiple comparisons (r=0.116, p= 0.068). Inclusion of the use of psychotropic medications (antipsychotics, anxiolytics, antidepressants, and sleep medications) into the model did not significantly alter these results. Baseline whole brain WMH load did not predict any other NPS severity scores.

**Figure 1.**
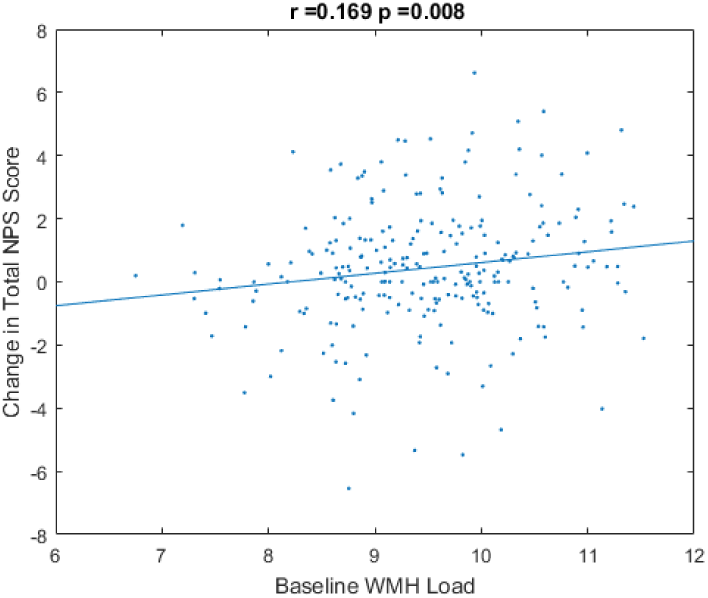
Correlation plot of baseline whole brain WMH load to predict the change in future NPI total scores over time

**Figure 2.**
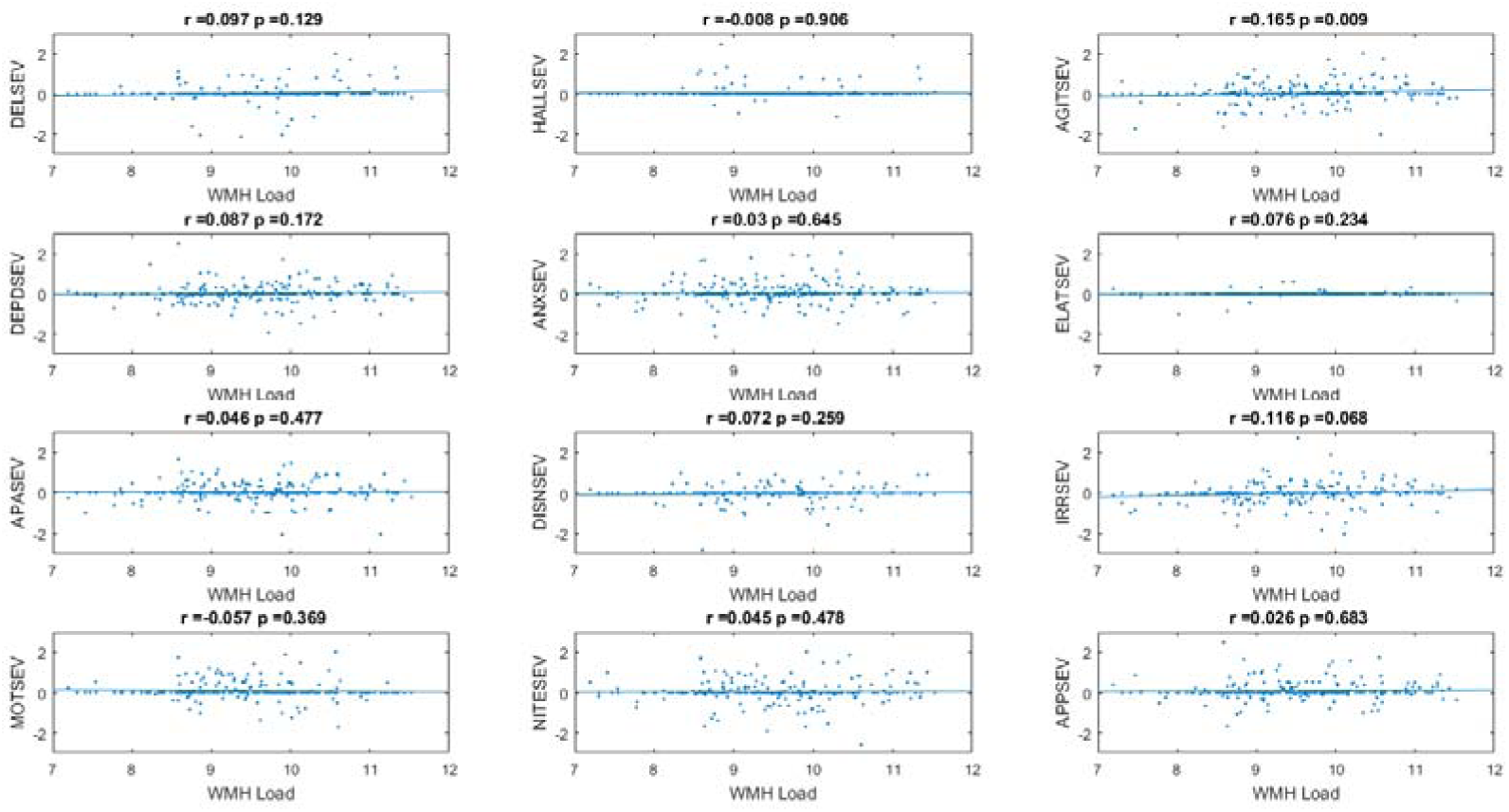
Correlation plots of baseline whole brain WMH load to predict the annualized change in future NPI severity scores over time, correcting for age, sex and education (without meds). From left to right: DELSEV = delusion severity score, HALLSEV = hallucination severity score, AGITSEV = agitation severity score, DEPD = depression severity score, ANXSEV = anxiety severity score, ELATSEV = elation severity score, APASEV = apathy severity score, DISNSEV = disinhibition severity score, IRRSEV = irritability severity score, MOTSEV = aberrant motor behaviour severity score, NITESEV = Night-time behaviour severity score, APPSEV = appetite severity score

As a secondary analysis, we determined which lobar WMH loads contributed most to the significant correlation between baseline whole brain WMH load and severity of agitation. The temporal lobe WMH (r=0.169, p=0.008) and frontal lobe WMH load contributed most to this finding (r=0.153, p=0.016) (Figure 3).

**Figure 3.**
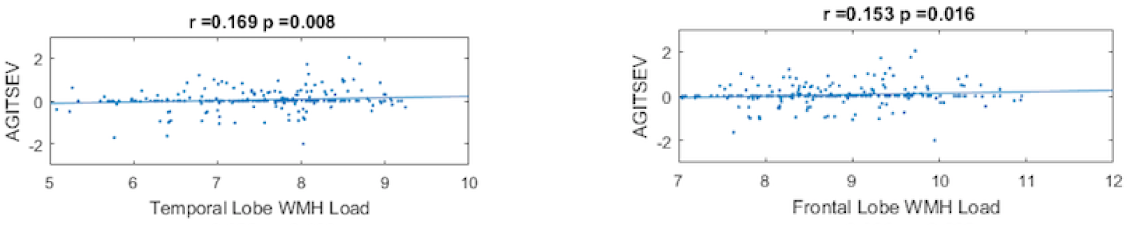
Correlation plots of baseline temporal and frontal lobe WMH load to predict the change in future agitation severity scores over time. AGITSEV = agitation severity score

## 4. DISCUSSION

The current literature regarding the relationship between WMH and NPS in patients with MCI or AD is inconclusive. As such in this study, we sought to assess the association between WMH load and NPS and found a significant relationship between baseline WMH and changes in the severity of NPS when conducting a longitudinal analysis in patients with MCI-AD and AD. Overall, our findings suggest that cerebrovascular injury in MCI-AD and AD, as measured by WMH load, is associated with the progression or worsening of certain NPS. Like others, our cross sectional results revealed no association between WMH and NPS(Klugman et al., 2009; Modrego et al., 2008; Rosenberg et al., 2015; Staekenborg et al., 2008). One previous study had divided cerebrovascular disease severity into four categories based on neuroradiology reports in an AD cohort and no association was found between WMH and NPS(Klugman et al., 2009). In another study, WMH were rated using the Wahlund scale and again no association between WMH and NPS was found in AD(Modrego et al., 2008). In a review on neuroimaging studies of agitation in AD(Rosenberg et al., 2015) only one cross sectional study examined the association between WMH load and NPS(Staekenborg et al., 2008) and they also found no association between WMH and NPS in AD. However, the WMH were not quantified but categorized according to the Fazekas scale. In all of these aforementioned studies where no association was found between WMH and NPS, WMH were not quantified but instead rated using visual scales. In contrast, a cross-sectional study using regional WMH volumes revealed that higher right frontal lobe WMH volume was associated with the presence of delusions in AD(C.J. Anor et al., 2017). In our current study, through post-hoc correlation analysis, we also found both baseline frontal WMH load and baseline temporal WMH load were correlated with the agitation severity score. The frontal and temporal lobes are often implicated in NPS(Boublay et al., 2016) and so it is not surprising that cerebrovascular disease in these areas would be an important contributor to NPS.

Although the regional (frontal lobe r=0.153, temporal lobe r=0.169) and whole brain (r=0.165) WMH loads effects on agitation are not large, it remains an important finding as WMH load reflects cerebrovascular disease, which is often due to modifiable risk factors such as hypertension and type 2 diabetes.In a recent study, pre-diabetes and diabetes have been shown to accelerate cognitive decline and predict micro-vascular lesions in healthy older adults(Marseglia et al., 2018). However, the effects of type 2 diabetes and other vascular risk factors mentioned above, can be significantly reduced by lifestyle changes. Modifications in diet and increased physical activity can help control vascular risk factors and reduce the risk of cardiac events(Aggarwal et al., 2018), which can contribute to the development of certain NPS.

In the longitudinal mixed effect model, baseline WMH load was associated with significantly worse future NPI total scores (p=0.014), NPI irritability severity subscores (p=0.009), and there was a trend for worse future agitation (p=0.058). Our finding that the baseline whole brain WMH load was associated with increased future irritability severity scores supports previous findings(Boublay et al., 2016; Ogama et al., 2018; Misquitta et al, 2019 under review). One study found focal frontal WMH to be associated with verbal aggressiveness in amnestic MCI and AD(Ogama et al., 2018), which supports our current finding of the association between frontal lobe WMH load and worsening agitation and irritability. Additionally, this finding echoes the results of Misquitta and colleagues (2019 manuscript under review), wherein they found WMH load to be a significant contributor to the NPI sub-syndrome of hyperactivity, which includes the following: agitation, euphoria, disinhibition, irritability, and aberrant motor behaviour. The current study suggests that irritability and agitation are the NPS within the sub-syndrome of hyperactivity that is associated with WMH load. These results are significant as both studies utilized large and established datasets: Alzheimer’s Disease Neuroimaging Initiative (ADNI) for Misquitta et al and NACC here.

Our study has a number of limitations. Although our sample was collected as part of a large standardized clinical data set, our final cohort was significantly decreased when excluding patients whose clinical data and MR imaging data were not within three months of each other, who did not have at least one follow up clinic visit, and who had a history of cognitive disorders other than AD. In addition, because the participants’ MRIs were collected with different MRI machines, this affected the quality of the images and increased the heterogeneity of images in our study. However, we implemented a previously validated automated WMH segmentation technique(M. Dadar et al., 2017) and utilized a library of manual segmentations based on the NACC dataset, verified by an expert (MD) to mitigate this issue. Therefore, the quality of the segmentations is reliable. Next, only baseline MRIs were used for this study, as we did not have enough follow up scans to match follow up NPI scores. Therefore, future studies investigating the relationship between NPS and WMH load in AD may benefit from having follow up MRI scans available for each participant in order to track simultaneously the change in WMH load and NPS. Finally, the baseline WMH are correlated with NPS sub-scores with small r-values, which only explains part of the variability. Other factors that could explain the variability are the effects of underlying AD pathology that lead to underlying structural and functional changes.

The contribution that WMH load has on NPS is likely a synergistic result of the underlying interactions between vascular and AD pathologies. More specifically, neurofibrillary tangles and amyloid-β are hallmarks of underlying AD pathology. These neurofibrillary tangles are found everywhere in grey matter and commonly found in subcortical regions, such as the hypothalamic nuclei, which is known to be involved in the regulation of certain NPS(Boublay et al., 2016). Therefore, the presence of WMH within these regions could cause further damage to that caused by the neurofibrillary tangles. One study also has postulated that SVD potentially could be involved earlier on in the AD process by hastening the accumulation of amyloid deposition due to improper perivascular drainage of amyloid-β(Ehrenberg et al., 2018). The interactions between vascular and AD neuropathologies, in addition to the significant role that genetics and environmental factors, may cumulatively contribute to NPS development and expression.

Overall, our study has shown a significant association between baseline WMH load and worsening of irritability and agitation over time. Given how common and distressful both NPS are to patients and their care providers and given that NPS can be clinical indicators of preclinical dementia syndromes, our findings have potentially important implications for treatment, arguing for aggressive treatment of cerebrovascular disease even in patients with MCI-AD and mild AD. By controlling and better managing vascular risk factors, it could reduce the burden of cerebrovascular disease and may also decrease the development of NPS in patients with MCI-AD and AD as a result.

## List of abbreviations

AD: Alzheimer’s disease
DLB: Dementia with Lewy Bodies
FTD: Frontotemporal dementia
MCI: Mild cognitive impairment
MRI: magnetic resonance imaging
NACC: National Alzheimer’s Coordinating Center
NPI: Neuropsychiatric Inventory
NPS: neuropsychiatric symptoms
VaD: Vascular dementia
WMH: white matter hyperintensities

## 7. ACKNOWLEDGMENTS

The NACC database is funded by NIA/NIH Grant U01 AG016976. NACC data are contributed by the NIA-funded ADCs: P30 AG019610 (PI Eric Reiman, MD), P30 AG013846 (PI Neil Kowall, MD), P50 AG008702 (PI Scott Small, MD), P50 AG025688 (PI Allan Levey, MD, PhD), P50 AG047266 (PI Todd Golde, MD, PhD), P30 AG010133 (PI Andrew Saykin, PsyD), P50 AG005146 (PI Marilyn Albert, PhD), P50 AG005134 (PI Bradley Hyman, MD, PhD), P50 AG016574 (PI Ronald Petersen, MD, PhD), P50 AG005138 (PI Mary Sano, PhD), P30 AG008051 (PI Thomas Wisniewski, MD), P30 AG013854 (PI M. Marsel Mesulam, MD), P30 AG008017 (PI Jeffrey Kaye, MD), P30 AG010161 (PI David Bennett, MD), P50 AG047366 (PI Victor Henderson, MD, MS), P30 AG010129 (PI Charles DeCarli, MD), P50 AG016573 (PI Frank LaFerla, PhD), P50 AG005131 (PI James Brewer, MD, PhD), P50 AG023501 (PI Bruce Miller, MD), P30 AG035982 (PI Russell Swerdlow, MD), P30 AG028383 (PI Linda Van Eldik, PhD), P30 AG053760 (PI Henry Paulson, MD, PhD), P30 AG010124 (PI John Trojanowski, MD, PhD), P50 AG005133 (PI Oscar Lopez, MD), P50 AG005142 (PI Helena Chui, MD), P30 AG012300 (PI Roger Rosenberg, MD), P30 AG049638 (PI Suzanne Craft, PhD), P50 AG005136 (PI Thomas Grabowski, MD), P50 AG033514 (PI Sanjay Asthana, MD, FRCP), P50 AG005681 (PI John Morris, MD), P50 AG047270 (PI Stephen Strittmatter, MD, PhD).

