## Supplemental tables 1 and 2 for "The Longitudinal Assessment of Neuropsychiatric Symptoms in Mild Cognitive Impairment and Alzheimer’s disease and their Association with White Matter Hyperintensities in the National Alzheimer’s Coordinating Center’s Uniform Data Set"

**SUPPLEMENTARY MATERIAL**

|  | Visit 1  (N=252) | Visit 2  (N=247) | Visit 3  (N=184) | Visit 4  (N=117) | Visit 5  (N=55) | Visit 6  (N=30) |
| --- | --- | --- | --- | --- | --- | --- |
| Average time between clinic visit and baseline MRI (years) | 0.06±0.1 | 1.06±0.3 | 2.23±0.5 | 3.26±0.5 | 4.37±0.8 | 5.61±1.0 |
| Average time between consecutive clinic visits (years) |  | 1.11±0.0 | 1.19±0.4 | 1.14±0.3 | 1.15±0.6 | 1.08±0.3 |

**Supplementary Table 1. The time frame (in years) between the baseline MRI and clinic visits, as well as the time frame (in years) between consecutive clinic visits (years).**

| **Delusions Severity Score** | | | | |
| --- | --- | --- | --- | --- |
|  | Estimate | SE | tStat | pValue |
| Age | 0.029 | 0.056 | 0.524 | 0.601 |
| Sex | 0.113 | 0.105 | 1.074 | 0.284 |
| Education | -0.084 | 0.051 | -1.642 | 0.102 |
| WMH | 0.022 | 0.055 | 0.393 | 0.694 |
| **Hallucinations Severity Score** | | | | |
|  | Estimate | SE | tStat | pValue |
| Age | -0.045 | 0.032 | -1.382 | 0.168 |
| Sex | 0.055 | 0.061 | 0.900 | 0.369 |
| Education | 0.170 | 0.030 | 5.714 | 0.000** |
| WMH | 0.009 | 0.032 | 0.289 | 0.773 |
| **Agitation Severity Score** | | | | |
|  | Estimate | SE | tStat | pValue |
| Age | -0.001 | 0.068 | -0.009 | 0.993 |
| Sex | 0.107 | 0.128 | 0.835 | 0.405 |
| Education | -0.102 | 0.063 | -1.638 | 0.103 |
| WMH | -0.051 | 0.067 | -0.756 | 0.450 |
| **Depression Severity Score** | | | | |
|  | Estimate | SE | tStat | pValue |
| Age | -0.141 | 0.063 | -2.222 | 0.027 |
| Sex | 0.199 | 0.119 | 1.665 | 0.097 |
| Education | -0.082 | 0.058 | -1.401 | 0.163 |
| WMH | 0.087 | 0.062 | 1.385 | 0.167 |
| **Anxiety Severity Score** | | | | |
|  | Estimate | SE | tStat | pValue |
| Age | -0.112 | 0.068 | -1.649 | 0.101 |
| Sex | 0.140 | 0.127 | 1.099 | 0.273 |
| Education | -0.034 | 0.062 | -0.546 | 0.586 |
| WMH | -0.020 | 0.067 | -0.295 | 0.769 |
| **Elation Severity Score** | | | | |
|  | Estimate | SE | tStat | pValue |
| Age | -0.035 | 0.053 | -0.656 | 0.513 |
| Sex | -0.049 | 0.100 | -0.484 | 0.629 |
| Education | -0.030 | 0.049 | -0.619 | 0.537 |
| WMH | -0.059 | 0.053 | -1.133 | 0.259 |
| **Apathy Severity Score** | | | | |
|  | Estimate | SE | tStat | pValue |
| Age | -0.029 | 0.063 | -0.454 | 0.650 |
| Sex | -0.097 | 0.119 | -0.816 | 0.415 |
| Education | -0.024 | 0.058 | -0.415 | 0.679 |
| WMH | -0.035 | 0.062 | -0.563 | 0.574 |
| **Disinhibition Severity Score** | | | | |
|  | Estimate | SE | tStat | pValue |
| Age | -0.026 | 0.064 | -0.411 | 0.681 |
| Sex | -0.046 | 0.120 | -0.385 | 0.701 |
| Education | -0.060 | 0.059 | -1.021 | 0.308 |
| WMH | 0.064 | 0.063 | 1.020 | 0.309 |
| **Irritability Severity Score** | | | | |
|  | Estimate | SE | tStat | pValue |
| Age | -0.057 | 0.066 | -0.860 | 0.391 |
| Sex | -0.180 | 0.125 | -1.443 | 0.150 |
| Education | -0.109 | 0.061 | -1.782 | 0.076 |
| WMH | -0.024 | 0.065 | -0.360 | 0.719 |
| **Aberrant Motor Behaviour Severity Score** | | | | |
|  | Estimate | SE | tStat | pValue |
| Age | 0.005 | 0.062 | 0.076 | 0.939 |
| Sex | 0.138 | 0.116 | 1.186 | 0.237 |
| Education | -0.021 | 0.057 | -0.376 | 0.707 |
| WMH | 0.034 | 0.061 | 0.550 | 0.583 |
| **Night-time Behaviour Severity Score** | | | | |
|  | Estimate | SE | tStat | pValue |
| Age | 0.068 | 0.072 | 0.951 | 0.342 |
| Sex | 0.116 | 0.135 | 0.863 | 0.389 |
| Education | -0.088 | 0.066 | -1.338 | 0.182 |
| WMH | 0.055 | 0.071 | 0.777 | 0.438 |
| **Appetite Severity Score** | | | | |
|  | Estimate | SE | tStat | pValue |
| Age | -0.057 | 0.061 | -0.936 | 0.350 |
| Sex | -0.056 | 0.115 | -0.486 | 0.628 |
| Education | -0.006 | 0.056 | -0.111 | 0.912 |
| WMH | -0.001 | 0.060 | -0.020 | 0.984 |

**Supplementary Table 2. Baseline mixed-effect model**
